# Supergene formation: evidence for recombination suppression among multiple functional loci within inversions

**DOI:** 10.1101/2021.12.16.472922

**Authors:** Paul Jay, Manon Leroy, Yann Le Poul, Annabel Whibley, Monica Arias, Mathieu Chouteau, Mathieu Joron

## Abstract

Supergenes are genetic architectures associated with discrete and concerted variation in multiple traits. It has long been suggested that supergenes control these complex polymorphisms by suppressing recombination between set of coadapted genes. However, because recombination suppression hinders the dissociation of the individual effects of genes within supergenes, there is still little evidence that supergenes evolve by tightening linkage between coadapted genes. Here, combining an landmark-free phenotyping algorithm with multivariate genome wide association studies, we dissected the genetic basis of wing pattern variation in the butterfly *Heliconius numata*. We showed that the supergene controlling the striking wing-pattern polymorphism displayed by this species contains many independent loci associated with different features of wing patterns. The three chromosomal inversions of this supergene suppress recombination between these loci, supporting the hypothesis that they may have evolved because they captured beneficial combinations of alleles. Some of these loci are associated with colour variations only in morphs controlled by inversions, indicating that they were recruited after the formation of these inversions. Our study shows that supergenes and clusters of adaptive loci in general may form via the evolution of chromosomal rearrangements suppressing recombination between co-adapted loci but also via the subsequent recruitment of linked adaptive mutations.

## Introduction

Recombination is a central force in evolution, allowing the reshuffling of genetic diversity and providing new combinations of alleles for natural selection to act on. Recombination is however also a homogenising force, breaking apart beneficial combinations of alleles and preventing the maintenance of alternative combinations. This is notably true when recombination occurs among combinations of alleles evolving under distinct selective pressures, for instance when populations evolving in heterogeneous environments are connected by gene flow. When adaptation in a given environment involves changes at multiple loci, heterogeneity in the environment generates selective pressure favouring either lower or higher recombination rates among these loci, depending on the strength and direction of selection that these loci experience (*1*) This is expected to lead to genomic variation in recombination rate, between and within chromosomes, and notably to the formation of clusters of locally adaptive loci (*1, 2*). Supergenes stand as notable instances of such clusters of adaptive loci, at which different combination of alleles are maintained within a single population.

Clusters of divergent loci and supergenes are observed in a broad diversity of organisms [e.g. (*3–5*)] but their origins is a long standing debate in evolutionary genetics (*2, 6–10*). Indeed, many processes, such as genetic drift or linked selection around a single selected locus, may lead to the formation of divergent clusters. Clusters involving several adaptive loci may evolve because of chromosomal translocation bringing unlinked loci into physical linkage or because those loci are captured by local chromosomal rearrangements suppressing recombination when heterozygous, such as inversions. Since new beneficial mutations are more likely to establish when they are linked to other variants under positive selection (a process known as divergence hitchhiking), clusters of adaptive loci and supergenes may also result from the serial recruitment by natural selection of mutations in tight linkage with other adaptive variants (*10, 11*).

Understanding the evolution of groups of tightly linked loci such as supergenes requires finely dissecting both the phenotypic effects of these loci across multiple related taxa and the origin of their linkage. Because of the scarcity of whole genome assemblies from groups of related species, as well as the complexity of untangling the genetic architecture of complex multidimensional traits, the evolution of clusters of adaptive loci, in particular supergenes, remains poorly understood. More critically, because recombination suppression causes loci captured by inversions to be in strong linkage disequilibrium, determining the individual effects of such loci has proven to be difficult (*12*). As a consequence, except for the specific cases of mating systems of plants (*13–16*) and fungi (*17, 18*), the individual contributions of loci maintained in linkage by supergenes remain largely unknown (*12*). This constitutes a major obstacle to our understanding of the evolution of supergenes and of genomes in general.

Neotropical butterflies in the genus *Heliconius* have been extensively studied over the past decades both ecologically and genetically. Most *Heliconius* species display geographic variations in wing pattern, involved in local mimicry with other butterfly species, and at least four major chromosomal regions are known to underlie these variations (*19–23*). Nevertheless, the genetic architecture of wing patterning seems to vary between species, as colour variations in most species involve only a subset of these four loci (*24, 25*). Because of this oligogenic control of mimetic colour pattern, crosses between different mimetic forms often result in the formation of recombinant, non-mimetic phenotypes. This is notably observed in the transition zones between the geographic ranges of distinct mimetic forms (*22, 26*) and the recurrent formation of these poorly protected individuals is expected to favor the evolution of clusters of wing pattern loci (*2, 27*). Consistent with this prediction, two independent inversions have been observed around *cortex*–a gene involved in wing patterning– in *H. numata* and *H. pardalinus* (*28*) and in *H. sara, H. demeter, H. hecalesia and H. telesiphe*, respectively (*29*). The sharing of inversions by multiple species was shown to result from multiple introgressions among closely-related taxa (*28, 29*).

Among these taxa, *Heliconius numata* appears as an outlier. Indeed, besides its geographical wing pattern variation, *H. numata* also displays a striking polymorphism of colour patterns within populations. This diversity is associated with three polymorphic inversions forming a supergene on chromosome 15 (Figure 1). In addition to the previously mentioned inversion called P_1_ capturing the gene *cortex*, this supergene also includes two other polymorphic inversions, P_2_ and P_3_, all in adjacent positions, and suppressing recombination over a 3Mb region encompassing 107 genes (Figure 1A) (*30*). This supergene, P, displays three allelic classes: Hn0, without any inversion, Hn1, with only the P_1_ inversion, and Hn123, with the three inversions P_1_, P_2_ and P_3_ (Figure 1A). Because the *H. numata* P supergene spans a region repeatedly found to be associated with wing pattern variations in Lepidoptera (*19, 24, 31*), it have been assumed that P inversions have evolved because they reduce recombination between several colour pattern loci and hamper the formation of maladapted recombinant phenotypes (*31, 32*), but this remained hypothetical.

**Figure 1.**
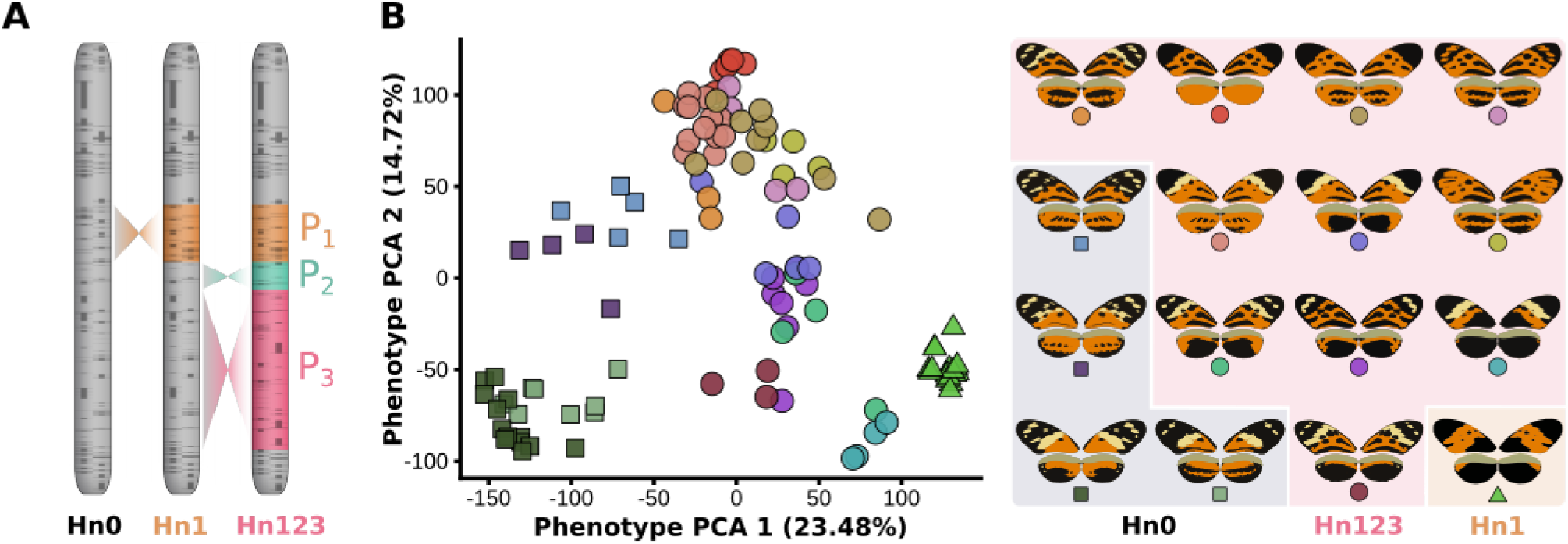
Genetic architecture and wing pattern diversity in *H. numata*. **A,** Genetic architecture of the *H. numata* mimicry supergene P characterized by three polymorphic inversions of respective size 400kb, 200kb and 1200 kb. **B,** Two-dimension approximation of the morphospace representing the phenotype diversity observed in *H. numata*. The dotplot display results from a Principal Component Analysis (the first two components are displayed here) computed on wing pattern variations as obtained from CPM (*33*). For display purpose, butterflies were manually classified into morphs based on litterature (*58*); different morphs are depicted by different colours. The butterflies sampled for this study represent the most common morphs observed in *H. numata*. Different supergene genotypes are depicted by different symbol shapes (square: Hn0/Hn0; circle: Hn123/Hn0, Hn123/Hn123; triangle: Hn1/Hn1, Hn1/Hn0, Hn1/Hn123). Results for PC 3 and PC 4 are presented in Figure S2. PCA was computed on samples belonging to the same supergene genotype in Figures. S3-4.

Here, to locate the loci underlying the phenotypic variation and forming this supergene, we took advantage of the fact that several morphs of *H. numata* share the same chromosomal arrangement at the colour-pattern supergene (Figure 1), and should therefore recombine normally, We re-sequenced the whole genomes of 130 specimens, used an unsupervised landmark-free algorithm to dissect their multidimensional wing-pattern variations and performed multivariate genome wide association studies to link phenotypic and genetic variations. We show that at least 19 genomic intervals are associated with colour variations in *H. numata* and most of them are in non-coding regions. All these regions are situated within the P supergene, bringing evidence that P inversions were recruited because of their role in maintaining beneficial combinations of wing pattern alleles. However, several of these regions seem not to be involved in wing pattern variations in closely related species and in *H. numata* individuals without inversions. This suggests that their involvement in *H. numata* colour variation evolved after the formation of the inversions, likely because their tight linkage with other wing pattern loci made them more prone to be recruited by natural selection than loci elsewhere in the genome.

## Results

In order to study the evolutionary events which led to the formation of the supergene P, we resequenced at approx. 30x coverage 131 *H. numata* classified following the litterature into 16 mimetic morphs (Figure 1) and mapped reads against the *H. melpomene* reference genome (Hmel 2). We used PCA to genotype for inversions [Figure S1, (*30*)], and found that 38 samples were homozygous for the ancestral gene order (Hn0/Hn0), 20 were homozygous or heterozygous for the first class of derived haplotype (Hn1/Hn1, Hn1/Hn0) and 73 samples were homozygous or heterozygous for the second class of derived haplotype (Hn123/Hn123, Hn123/Hn0, Hn123/Hn1).

Morphometric analyses were run on 110 samples whose wings were in good condition using Color Pattern Modelling (CPM), an algorithm for the quantification of colour pattern variations based on color classification and color pattern registration (*33*). Briefly, starting from standardized pictures, wings were first extracted, their colours clustered into black, orange or yellow (the three colours present on the wings), and then aligned with each other on the basis of pattern similarity. Each pixel common to all aligned wings was considered a variable, resulting in the description of wing pattern variation by ca. 10^5^ variables. Principal component analyses (PCA) were used to reduce the high dimensionality of the colour variation, and, as expected, showed that wing pattern polymorphism involves a mixture of qualitative and quantitative variation (Figures 1B and S2-4). Wing pattern polymorphism in *H. numata* indeed involves variations on different parts of the wing, with some features appearing more discrete (e.g. presence of a broad yellow band) than others (e.g. spread of hindwing black patterns). Supergene inversions appear as major determinants of specimen phenotype since samples with similar supergene genotype are phenotypically clustered (Figure 1B). By contrast, no wing pattern feature is associated with a specific geographic locality and the origin of samples appears as a poor descriptor of individuals phenotype (Figure S5). Moreover, we found no genetic structure in *H. numata* across the Neotropics (Figure S6; see also (*34*)), with the exception of samples from the Brazilian Atlantic forest which were removed for subsequent analyses. In total, 102 phenotyped and genotyped samples were retained in the analysis.

In order to identify the loci associated with wing pattern variation, we performed with MV-PLINK (*35*) multivariate Genotype-Phenotype associations using as phenotype the first six principal components describing the joint variation of fore and hind wings (variance explained: 58.08%). As expected following previous studies (*19, 30*), the main region of association corresponds to the P supergene (Figure 2A). A peak of association on chromosome 7 was found to be caused by an assembly error in the *H. melpomene* Hmel2 reference genome and this region actually maps to chromosome 15, in the supergene region (see Methods). The P locus therefore appears as the main determinant of phenotype variation in this sample set. Because inversions involve segments in opposite orientation that do not recombine, many mutations in the supergene were found to be in strong linkage disequilibrium and therefore equally associated with phenotypic variation (Figure 2B). To remove the confounding effect of inversions, we performed phenotype-genotype associations separately on the samples belonging to allelic classes Hn0 and Hn123 (Hn1 specimens were not analysed since they are all phenotypically very similar; Figure 1B).

**Figure 2.**
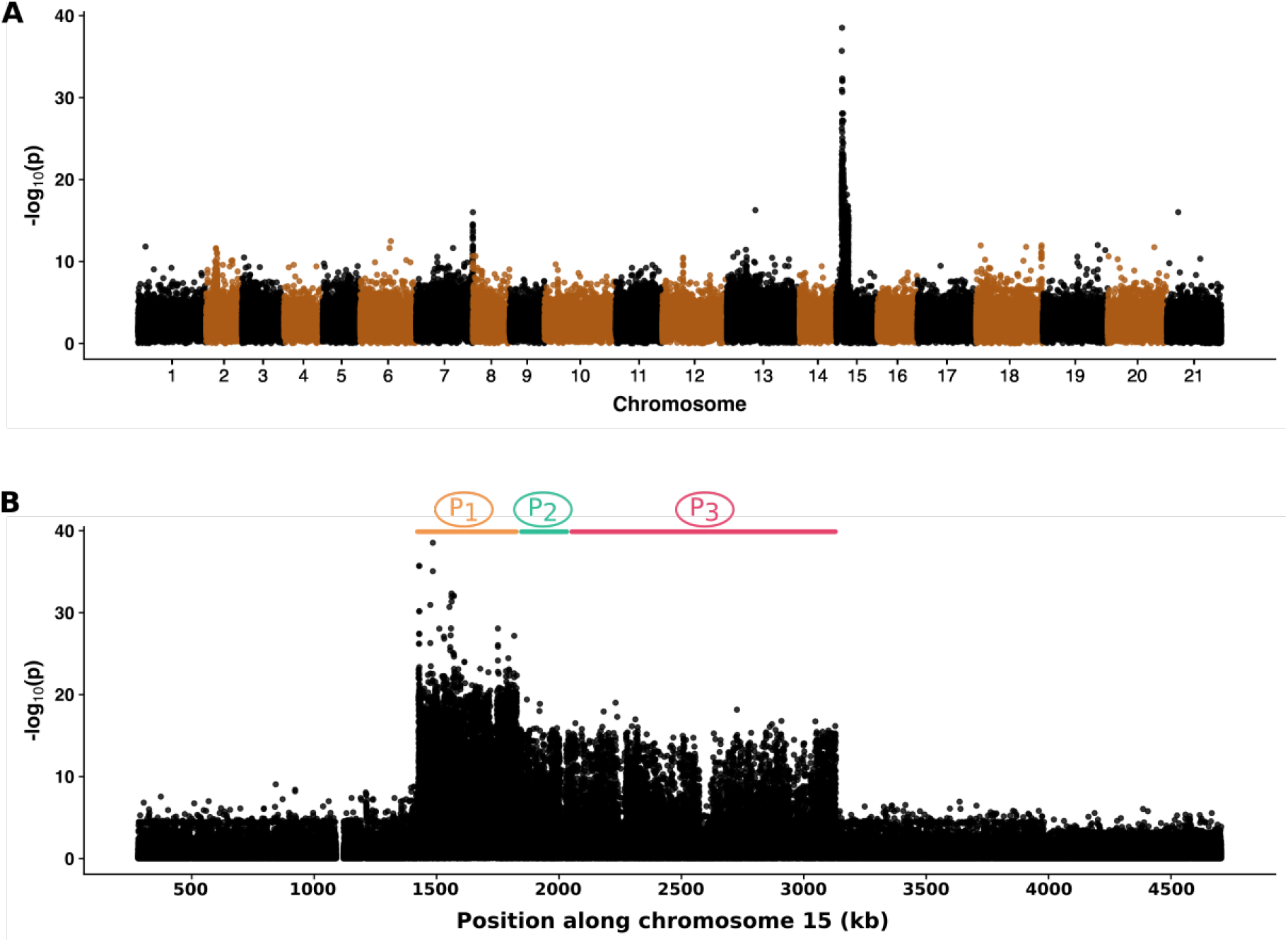
Genome wide association study with wing pattern variations. **A,** Multivariate association study using as phenotype the first six principal components describing wing pattern variations. One major peak of association in noticeable on chromosome 15 corresponding to the supergene. One minor peak can be seen on chromosome 7. This is due to an assembly error, and in reality this region is within the supergene region (see methods). **B** Focus on the peak of association on chromosome 15, corresponding to the position of the three polymorphic chromosomal inversions P_1_, P_2_ and P_3_.

In accordance with the model of a supergene composed of multiple genetic elements controlling individual trait variation, we expected to observe different regions of the supergene associated with different features of the wings. Genotype-phenotype associations performed separately on the two allelic classes indeed showed that different regions within the supergene are associated with wing pattern variations (Figure 3A and S7). When computing analyses using variation of the entire wing pattern, we observed that several regions of the supergene displayed very close peaks of association, making it difficult to distinguish them from each other. To overcome this limitation, we computed analyses focusing on more specific components of wing pattern variation (Figure 3A), such as the colour variation found on hindwings only, or the variation in yellow patterning, for example. Such analyses revealed well defined, sizeable peaks of association with different features of phenotype (Figure 3A, Figure 4 and S7). Manually comparing the results obtained with these analyses, we identified 19 regions with well-supported association with wing pattern variation in the Hn123 allelic class (Figure 3A; Table S2), in Hn0 (Figure S7; Table S2) or in both (Table S2). Among the most associated variants in each of these 19 regions (three variants per regions were considered), only one sits in a exon and is a non-synonymous mutation, i.e. these regions fall primarily in noncoding intergenic or intronic sequences (Table S2).

**Figure 3.**
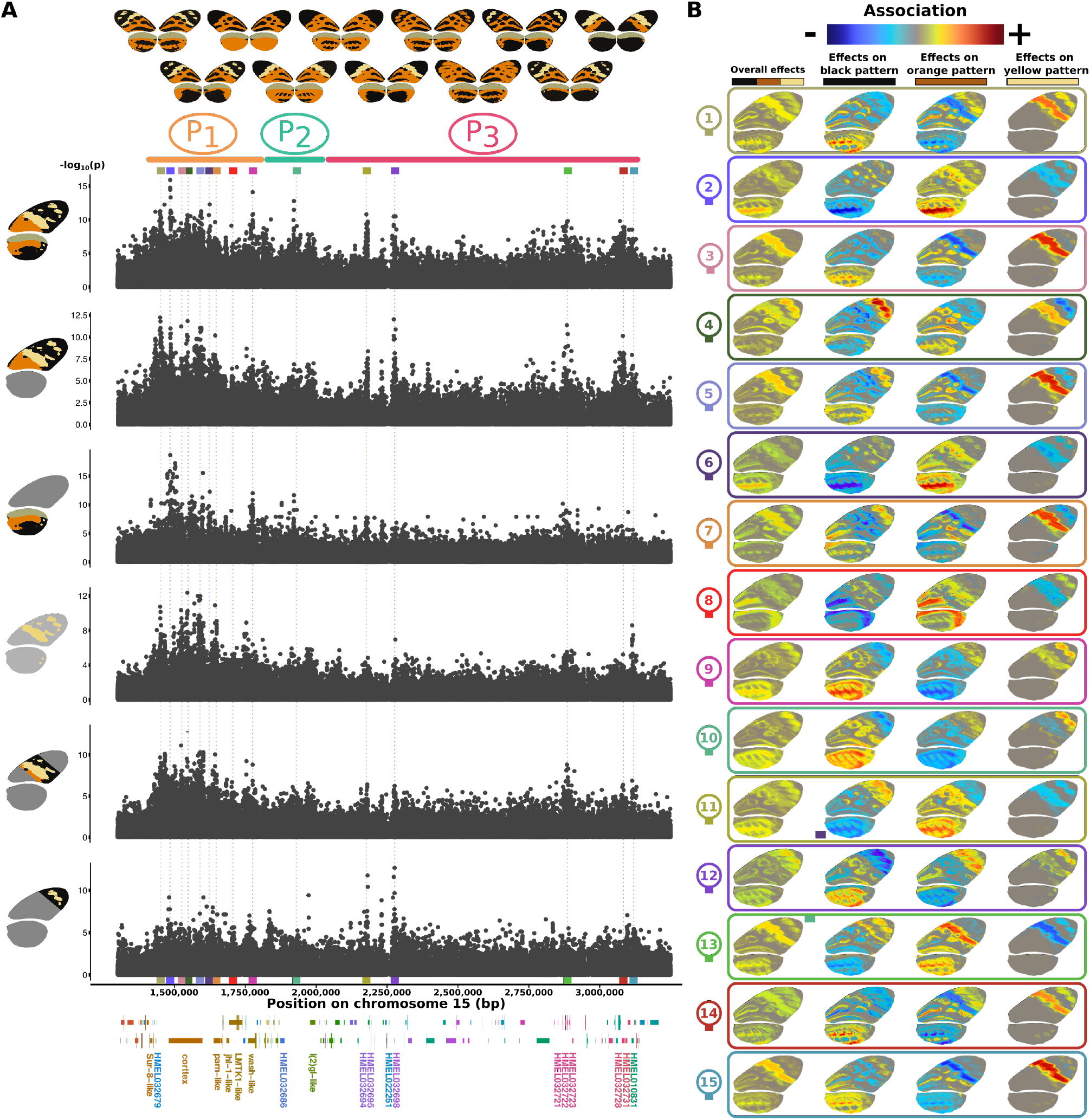
Distinct regions within the supergene are associated with variation of wing pattern features. **A**, Multivariate association studies computed on morphs controlled by the Hn123 arrangement, for different partitions of the wing: hindwing and forewing together, forewing, hindwing, yellow pattern on both wings, middle part of forewing, and tip of forewing. Coloured rectangles below plots indicate regions with association peaks [i.e. several variants with p<1e^−7^] in at least one of the partitions studied (Table S2). Data for the Hn0 arrangement are presented in Figure S7. **B**, Phenotypic effects of the top variant from the each of the 15 regions with high association to the wing pattern in Hn123 arrangement (coloured rectangles). Heatmaps from blue to red represents the strength and the direction of association of the derived allele for every pixel, that is how the allelic change at a given genetic position affect this wing area. Overall effects are shown as well as colour-specific effects, this latter representing to what extent the allelic change is associated with the presence or absence of this colour at this wing area. Because blue and red represent the direction of the association, opposite direction (i.e. red and blue values) on the same wing area when comparing two colour-specific heatmaps indicates that the focal locus is associated with a change from one colour to the other at this wing area. For instance, if the effect of a genetic variant on a given wing area is highlighted in blue when looking at orange patterns but in red when looking at black patterns, it means that this variant is associated with a switch from orange to black at this wing area.

**Figure 4.**
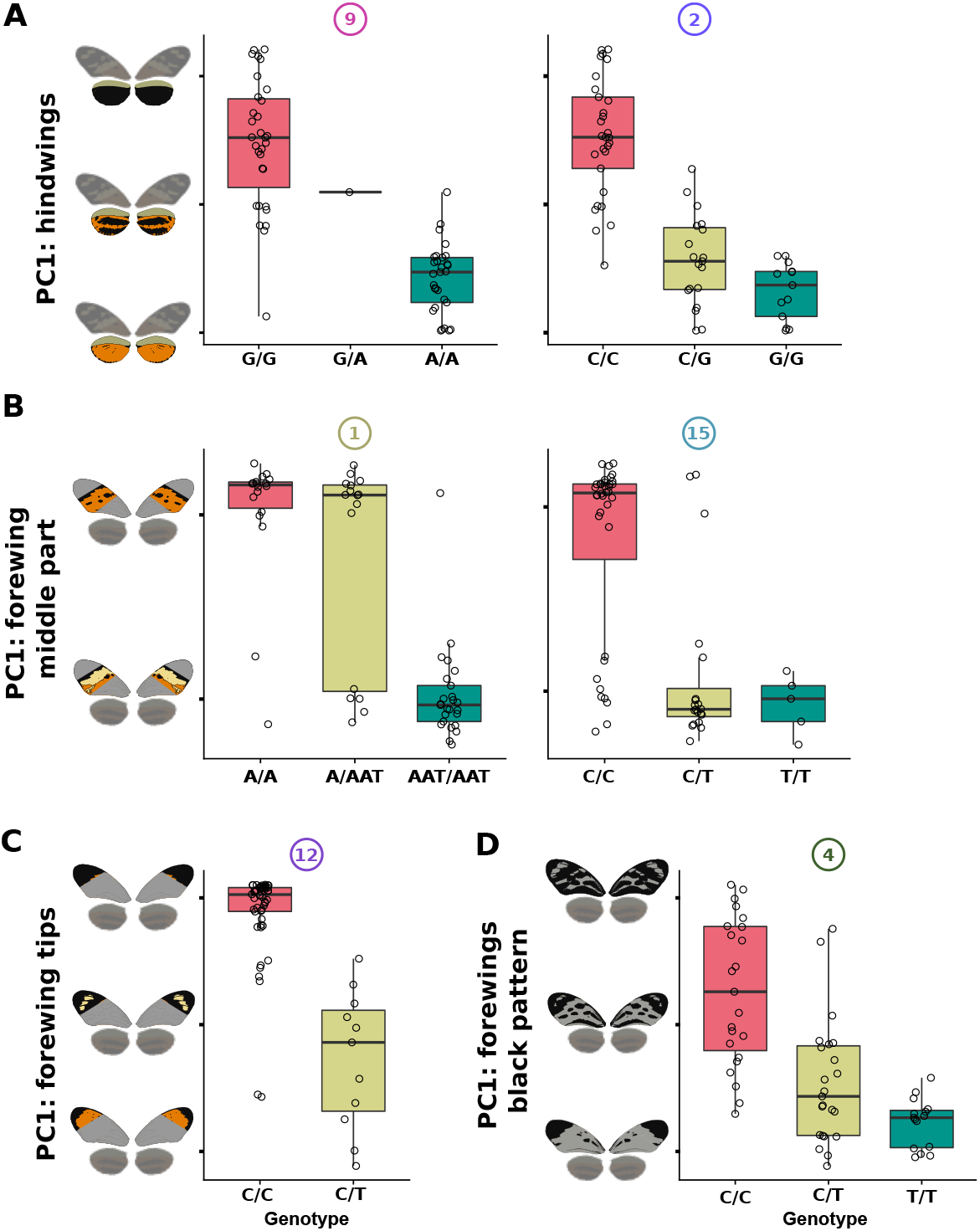
Effect of six selected variants on Hn123 wing pattern variations. The first principal components of analyses computed on different parts of the wing are used as proxy of the phenotype (y-axis): PCA computed on hindwings only (A), on the middle part of the forewings only (B), on the tips of the forewings only (C), black patterns of the forewings (D). Each dot is an individual. Instead of annotating y-axis with eigenvalues (values of individual on principal components), we display schematic butterflies with average phenotypes along the principal components. Boxplot elements: central line, median; box limits, 25th and 75^th^ percentiles; whiskers, 1.5 × interquartile range. **A**, Effect of the most strongly associated variants in regions 2 (*Cortex*, intron 2) and 9 (*Wash*, intron 2) on the amount of black on hindwings. **B**, Effect of the most strongly associated variants in region 1 (intergenic *HMEL032679-Cortex*) and 15 (intergenic *HMEL032731-HMEL010831*) on the presence of a yellow band on forewings. **C**, Effect of the most strongly associated variant in region 12 (intergenic *HMEL022251-HMEL032698*) on the tip of the forewings. **D**, Effect of the most strongly associated variant in region 4 (*Cortex*, intron 23) on the amount of black in the forewings.

In order to determine in more detail the wing pattern features affected by each genomic region, we computed the effect of the most associated variant (SNP, insertion or deletion) of each of these regions on the colour variation at each wing position (image pixel). This can be visualised in the form of heatmaps of variant effects on the wings, where colour hue and brightness reflect the direction and the strength of effect. Because each pixel can take three different colours (black, yellow and orange), the effect of variants on the presence or absence of these colours can be visualized for each colour independently but can also be summarized into an overall, non-directional effect per pixel (Figure 3B and S7). Most variants showing a significant association [p<1e^−7^] with wing colour pattern were involved in the variation of multiple elements of wing colouration, such as, for instance, the spread of black patterning on hindwings and the presence of a yellow band on forewings. This may reflect a property of our dataset (correlation of different wing pattern features in the forms used in our analysis), or may truly reflect pleiotropic effects of the mutations. Interestingly, closely-linked regions of the same gene were sometimes observed to correlate with different features of wing patterns. For instance, some regions of the gene *cortex* were associated with change in yellow features of the forewing whereas some very close regions of the same gene were associated with change in black features in hindwing (Figure 3 & 4, Table S2). Even though the limited phenotype variation among Hn0 individuals did not enable proper dissection of the variation of the different wing pattern features for those samples (Figure S7), several regions were associated with phenotype changes in samples with (Hn123) and without inversions (Hn0, Figure 7 and S7 and Table S2). For instance, the same intron in the gene *parn* is associated with colour variation in Hn0 and Hn123 samples.

In Hn123 specimens, we were able to finely dissect the genetic architecture underlying various features of the wing pattern, for instance locating regions respectively associated with presence of yellow band on the middle of the forewing, with yellow patches on the tip of the forewing and with the variation in size of black patches on hindwings (Figure 3 & 4, Table S2). All the aspects of the wing pattern variation studied here showed an association with multiple genomic regions of the supergene. The co-association of different regions with the same traits may result from an absence of recombination between these regions or because natural selection foster their coupling (for instance because these loci are epistatic). Genome assembly comparisons (*30*) and patterns of linkage disequilibrium within each allelic class (Figure S8-9) do not suggest the existence of additional chromosomal rearrangements or other mechanisms suppressing recombination between these co-associated regions, meaning that their coupling is likely maintained by selection. In addition, different regions associated with similar phenotypic variations are characterized by different sets of heterozygous or homozygous specimens for ancestral and derived variants, confirming that these regions recombine within each allelic class (e.g. Figure 4A-D).

Reanalysing previously published RNAseq data (*30, 36*), we found that genes *Cortex, Parn, Wash*, lgl, *jhI-1* and *HMEL032728* display significant expression differences in wing disk when comparing allelic classes (Figure S10). Since these genes include or are very close to regions associated with wing pattern variation in our analyses (Figure 3, Figure S10, Table S2), their differential expression during wing development strongly suggests that they participate in the control of wing pattern variation in *H. numata*.

Phylogenomic and phenotypic analyses show that the genetic variants associated with some wing pattern features seem to have evolved after the formation of inversions. Some phenotypes and genetic variants are indeed only observed in a subset of the morphs belonging to the Hn123 allelic class (Figure 1B). Since the formation of an inversion is a unique event leading to the capture of a single haplotype (i.e. inversions do not include any polymorphism upon formation, (*30*)), variants restricted to a subset of Hn123 morphs have necessarily evolved after the formation of this allelic class and the occurrence of the inversions. For instance, the morphs *aurora, timaeus* and *tarapotensis* are all associated with the Hn123 supergene allele (with inversions P_1_, P_2_ and P_3_) but differ in several respects, such the presence of a median yellow band on forewings, or spots on forewing tips (Figure S11). Phylogenomic analyses using topology weighting show that, along the supergene, alleles controlling the morphs *tarapotensis* and *timaeus* tend to often cluster together relative to *aurora*. However, topologies shift repeatedly along the inversions and, at some positions associated with wing pattern variation, *timaeus* may tend to cluster with *aurora* (Figure S11). Taken together, these results suggest that the genotype-phenotype associations found at the P supergene do not only result from the capture of previously-segregating variants and their subsequent protection against recombination by inversions, but also from the evolution and diversification of these haplotypes after inversion formation.

## Discussion

In summary, within the supergene interval, multiple genomic regions are associated with different wing pattern features, lending support to the hypothesis that the supergene locks together multiple, independent sites each associated with specific features of the phenotype under selection. This therefore support the « supergene model » (*12*) or “beads-on-a-string” model (*37*), and is generally inconsistent with the hypothesis that the supergene could involve a single gene with pleiotropic effects (*38*). Since our findings are based on genotype-phenotype correlations, we cannot exclude that not all of these loci would be functionally involved in wing colouration. An association could result, for instance, from a correlation between wing pattern traits and certain unmeasured traits. In this species, morphs share the same microhabitat and display similar behaviour (*39*); however, they tend to mate disassortatively, i.e. females preferentially mate with males displaying a different wing pattern (*40*). Depending on how this preference is genetically determined (*41*), it might cause spurious genotype-phenotype association. For instance, if a locus controlling a given wing pattern feature is in linkage with a locus inducing avoidance for such feature during courtship, genetic mapping will associate both loci to wing pattern variation. Nonetheless, this is unlikely to result in multiple non-causal associations. Moreover, such associations would reveal coordination of wing pattern variants and mate choice variants, which still stresses the importance of chromosomal rearrangements in maintaining co-adapted loci in linkage.

Some of the genes in wing-pattern associated regions display RNA expression differences during wing development when comparing morphs, suggesting that these regions are indeed involved in wing patterning. Among these genes, *cortex, parn* and *wash* are also associated with wing pattern variations in other Heliconius species (*19, 24, 25, 42*), but only the role of *cortex* and *wash* have been experimentally validated. Taken together, these results indicate that several loci associated with wing pattern variations in the supergene interval of *H. numata* are most likely functionally involved in wing patterning. This confirms the long-standing assumption that the supergene P coordinates the variation of genetic elements underlying the distinct mimetic morphs in *H. numata*. Our result shows that *cortex* and *wash* controls clearly different features of wing pattern (*42*) and also highlight that several other loci linked to these genes may also play an important role in wing-patterning, such as the genes *parn, l(2)gl* and *jhI-1*. These genes regulate general processes such the transcription or cell division in insects, and *l(2)gl* mutants in Drosophila notably show aberrant wing development (*43*). They may therefore constitute good candidates for controlling wing colour pattern in Heliconius, but functional studies are required to better understand their role

Some of these loci captured by *H. numata* inversions are thus associated with wing pattern variations in *H. numata* specimens with and without inversions, and in other species. This strongly suggests that P inversions have evolved because, via their effect of suppressing recombination, they maintain beneficial combinations of wing pattern alleles at these loci, forming good mimetic morphs. Non-mimetic morphs in *H. numata* have indeed been shown to be strongly selected against by bird predation (*40, 44*), which constitutes a powerful selection pressure on mechanisms limiting the formation of recombinant morphs. The P_1_ inversion is also found in *H. pardalinus* (*28*) and recent studies have found two other inversions at a similar location, respectively in *H. sara, H. demeter, H. hecalesia* and *H. telesiphe* (*29*), and in a very distant swallowtail butterfly, *Papilio clytia* (*45*). The benefits of recombination suppression induced by chromosomal rearrangements have extensively been studied theoretically (e.g. (*2, 27*)), but empirical evidence is very scarce due to the complexity of performing genetic mapping of non-recombining regions. The convergent evolution of inversions at this region on chromosome 15 in Lepidoptera and the findings that this region includes multiple wing-pattern loci provide a rare empirical demonstration of the benefit of recombination suppression.

The supergene interval constitutes a dense cluster of co-adapted loci in a relatively small genomic region (3Mb). Synteny in the supergene interval is highly conserved across Heliconius and more broadly in the Lepidoptera (*29, 30, 46*), indicating that this cluster does not result from recent translocations of loci from other chromosomal regions. Some of the loci appear to be associated with wing pattern variation only in some *H. numata* morphs carrying inversions and not in samples without inversions (Hn0) or in other Lepidoptera. For instance, *H. ismenius*, the sister species of *H. numata* (2.54 Mya of divergence), includes several mimetic races phenotypically very similar to certain *H. numata* morphs, but chromosome 15 variations appear to affect only minor aspect of wing pattern, while most wing pattern variations are controlled by loci on chromosomes 18 and 10 (*25*). The involvement of chromosomes 18 and 10 in wing patterning are found in most Heliconius subclades, such as in *H. hecale, H. erato, H. cydno, H. melpomene*, but not in *H. numata* (*25*). The absence of association at these loci in *H. numata* and the existence of loci solely associated with wing pattern variation in *H. numata* indicate that different networks of genes controlling colouration have evolved during the divergence of these taxa. These putative new wing pattern loci within the supergene inversions may have resulted from their recruitment by natural selection because of their tight linkage with other wing pattern loci within these inversions,. Following theoretical predictions (*10, 11*), inversions could have evolved because they maintain in linkage loci initially involved in wing patterning, and the resulting reduced recombination over a sizeable region may have subsequently favoured the recruitment of additional adaptive mutations in this region (divergence hitch-hiking; (*47*)) thereby leading to the formation of new loci encoding wing pattern variation. This implies the pre-existence in the supergene interval of loci functionally involved in wing development, but previously undetected because of a lack of association with wing pattern variation in other taxa. Experimental assays are required to understand the implication of these loci in wing patterning across the Heliconius clade.

In summary, we found multiple loci associated with different wing pattern features in the *H. numata* supergene. Several of them are very close to genes differentially expressed in wing discs between morphs and are also associated with wing pattern variations in related taxa. We found no unusual recombination pattern that could cause spurious wing pattern associations, and these associations are not expected to result from the correlation between wing pattern features and unmeasured traits. Overall, it indicates that several of these loci may be indeed functionally involved in wing patterning. In accordance with theory (*2, 10, 11*), we found that the phenotypic diversity in *H. numata* is encoded by a tight cluster of loci, whose formation likely results both from chromosomal rearrangements suppressing recombination between alternative, beneficial combinations of alleles at coadapted loci, and from the further recruitment of new adaptive loci within these inversions, probably due to the increased recruitment probability of beneficial alleles when in linkage with other beneficial mutations. Our results provide empirical evidence that chromosome rearrangements evolve because they maintain linkage between coadapted loci, and, therefore, that supergenes allow the switch between combinations of coadapted alleles, which was long predicted but have received very few empirical demonstration (*12*). Moreover, we revealed that *H. numata* and its closely related sister species display radically different genetic control of similar phenotypic variations, showing that natural selection can favour the rapid evolution of alternative genetic architectures.

## Methods

### Sampling and sequencing

Resequenced genome data from (*30*) and (*28*) were used and were completed by 62 new specimens. In total, 131 specimens belonging to 16 mimetic variant of *H. numata* and from five geographical origins: Peru (n=85), Ecuador (n=13), Colombia (n=10), French Guaina (n=6), Brazil (n=17) were used (Table S1). Wings were preserved at room temperature and bodies were conserved in DMSO at −20 °C.

DNA was extracted from thorax tissue using the Qiagen DNeasy blood and tissue extraction kit. Illumina Truseq paired-end libraries were prepared and sequenced in 2×100 base pair on an Illumina NovaSeq platform (Get Plage, INRA Toulouse). Reads were mapped on the *H. melpomene* (Hmel2) reference genome (*48*) with NextGenMap (*49*) with default parameters. Mapped reads were processed with GATK and SNP detection was performed with the *unified genotyper*, following the procedure recommended by the authors (*50*). Bcftools was used to process the vcf files (*51*)

The misplacement of the scaffold on chromosome 7 (instead of on chromosome 15, see main text) was determined by mapping this scaffold with blast on NCBI database and by checking that this error was corrected in the new Heliconius reference genome (Hmel2.5, published after the performance of these analyses, (*52*))

### Population genetic analyses

Principal component analyses were computed on SNP data using SNPRelate (v.3.9; (*53*)). This was used to assess genotypes at the P supergene and to detect whole genome *H. numata* geographical structure. Fst scans were performed using scripts from https://github.com/simonhmartin/genomics_general, using 5000 bp windows with a least 500 SNP per window. Linkage desequilibrium analyses were computed with tomahawk (*https://github.com/mklarqvist/tomahawk*). Because populations from the Atlantic forest of Brazil were shown to be differentiated from other *H. numata* populations, we removed them from subsequent analyses. RNAseq data from (*36*) were reanalysed using the EdgeR R package^*63*^ (v3.16.5). Gene expression in early pupal (24h) wings discs from *silvana* individuals (Hn0/Hn0) was compared to gene expression in both *tarapotensis* and *aurora* individuals (Hn123/Hn123). In order to study the evolutionary history of the supergene, we computed sliding window phylogenies along the supergene, using 10 kb windows with RaxML (*54*). Only sample homozygous for the inversions or for their absence were used (Hn0/Hn0, Hn1/Hn1, Hn123/Hn123). Moreover, based on PCR genotyping (*30*) and breeding experiment results (data not shown), we removed samples that might be heterozygous for two supergene alleles belonging to the same allelic class. For instance, considering that different Hn123 haplotypes encode the morphs *aurora* and *tarapotensis*, respectively, a Hn123/Hn123 individuals might have an a*urora*/t*arapotensis* genotype, which could confound phylogenetic analysis studying the evolution of these two haplotypes. Because haplotypes of the same allelic class recombine and because there are likely many neutral polymorphisms segregating in each classes, phylogeny topology were highly variable along the supergene. To summarise these variations (topology weighting), we used Twisst (*55*) using the different morph as different taxa. We used the morphs *silvana, bicolouratus, aurora, timaeus, lyrcaeus, tarapotensis* and *messene* to perform such analyses *s*ince they were the only morphs with sufficient samples.

### Phenotyping

Colour pattern variation was described using Color Pattern Modeling (CPM) according to the developer’s recommendations (*33*). Briefly, from standardized images, CPM quantifies phenotypic variation among specimens by producing comprehensive descriptors of the colour pattern which avoids the user making assumptions about the relevance of certain descriptors. Wing photographs were colour-segmented automatically and the resulting colour partitions were attributed to one of the three colours composing the tiger patterns displayed by *H. numata* morphs (and their comimics): black, orange, and yellow. Wing were aligned to an average model (improved by recursion), by maximizing the mutual information between individual pattern and the model. Wingpattern phenotypic variation could then be described as the color variation among all common pixel to all aligned wings. This high dimensionality space (ca. 10^5^ pixels) was summarized by principal component analysis (PCA).

In order to isolate variants associated with specific features of the wing pattern, we also performed CPM analyses on different part of the wings. Besides the analysis of fore- and hindwings together, we analysed separately the following partitions: (ii) forewings, (iii) hindwings, (iv) black patterns of the forewings, (v) black patterns of the hindwings, (vi) yellow patterns of the forewings, (vii) base of the forewings, (viii) median area of the forewings (yellow band area), and (ix) apex of the forewings. To take into account the genetic structure of the supergene P, which implies the absence of recombination between samples harbouring different rearrangements, we performed PCAs on subset of samples, based on their genotype (presence/absence of the three inversions). To compute the effect of SNP on the colour variation at each wing position (image pixel), we translated the loads (contribution to the multivariate association result) attributed by MV-PLINKs to each phenotypic traits (i.e. to each phenotypic principal components) into pixel values using the eigenvalues of the phenotypic PCA. See MV-PLINK and CPM references for further details (*25, 33, 35*).

### Multivariate association studies

To determine the genetic basis of colour pattern variation, multivariate genome wide association studies were performed using MV-PLINK (v1.6; (*35*)). MV-PLINK performs a Canonical Correlation Analysis (CCA) to test for an association between variation at multiple phenotypes at once and a single SNP. Here, we used as phenotypes the principal components from the PCA computed on phenotypic variations (obtained with CPM). The description of variation in width, size and translation of wing patterns elements increases in accuracy with the number of principal components considered (*56*). Nevertheless, including many non-informative phenotypes (i.e. principal components) in multivariate associationx causes non-informative association results, which hampers isolating meaningful associations. Thus, we calculated G*E association using up to 10 principal components as phenotypes, and retained the analyses allowing the best visualization of well-define peak of association. To take into account the supergene structure, GWAS were carried out independently within each genotypic group Hn0 (Hn0/Hn0), Hn1 (Hn1/Hn1, Hn0/Hn1) and Hn123 (Hn123/Hn0, Hn123/Hn1, Hn123/Hn123). Only SNPs with minor allele frequency (MAF) > 0.02 and genotyping rate > 0.5 were conserved for analyses, resulting in 532 574 SNPs used for Hn0 association and 306 921 SNPs for Hn123 association at P region. SNPeff (v4.3; (*57*)) was used to annotate genetic variants based on *H. melpomene* reference genome annotation. Region of association were manually defined by identifying 10kb regions with distinguishable peak of association including SNPs with p<1e^−7^ (Table S2). However, we acknowledge that many regions present many highly associated SNP but not forming a clear peak. This likely result from the tight physical linkage of loci associated with wing pattern variation, especially around *cortex*. Hence, the list of associated regions and SNP presented in Table S2 is not intended to be exhaustive but reflects the regions that we think are of interest, because of clear peaks of association or because of their strong association with particular wing pattern features.

## Acknowledgment

We are very grateful to Oscar Puebla, Floriane Coulmance, and to the Puebla lab for their careful review and useful comments on this manuscript. We thank the Peruvian government for providing the necessary research permits (236-2012-AG-DGFFS-DGEFFS, 201-2013-MINAGRI-DGFFS/DGEFFS and 002-2015-SERFOR-DGGSPFFS). This research was supported by Agence Nationale de la Recherche (ANR) grants ANR-12-JSV7-0005 and ANR-18-CE02-0019-01 and European Research Council grant ERC-StG-243179 to MJ. This project benefited from the Montpellier Bioinformatics Biodiversity platform supported by the LabEx CeMEB, ANR “Investissements d’avenir” programme ANR-10-LABX-04-01.

## Data availability

Scripts used to produce analyses have been deposited on Github: https://github.com/PaulYannJay/HeliconiusGWAS. The raw sequence data were deposited in NCBI SRA and accession numbers are indicated in Supplementary table 1 (the new genomes will be deposited upon manuscript acceptance). The whole genome VCF file is available upon request.

## Author contributions

P.J. and M.J. designed the study; P.J. wrote the paper; P.J. and A.W. generated the genomic data; Butterflies were collected by P.J., M.J., M.C. and M.A.; P.J. and M.L. performed the association studies; P.J., M.L. and Y.L.P. performed phenotyping; All authors contributed to editing the manuscript; M.J. supervised the study.

## Notes

### Competing Interest Statement

The authors have declared no competing interest.

