## Supplementary material (Figures S1-11) for "Supergene formation: evidence for recombination suppression among multiple functional loci within inversions"

This document contains:

Fig. S1-11

Table S1-2

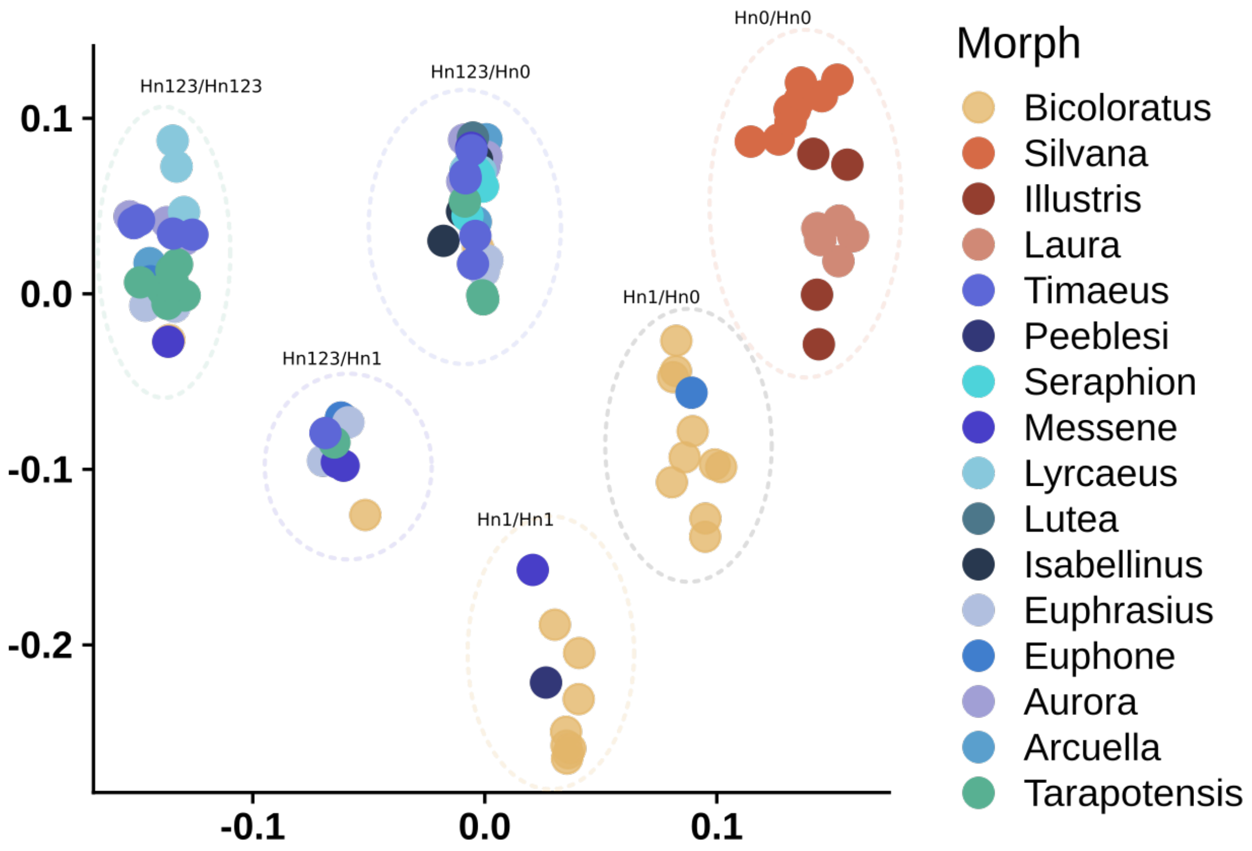


**Fig. S1 | Genetic structure observed at the supergene**

Principal component analysis computed on SNP data at the region on the supergene. Six clusters of samples can be observed, and correspond to the six distinct genotypes at the supergene.

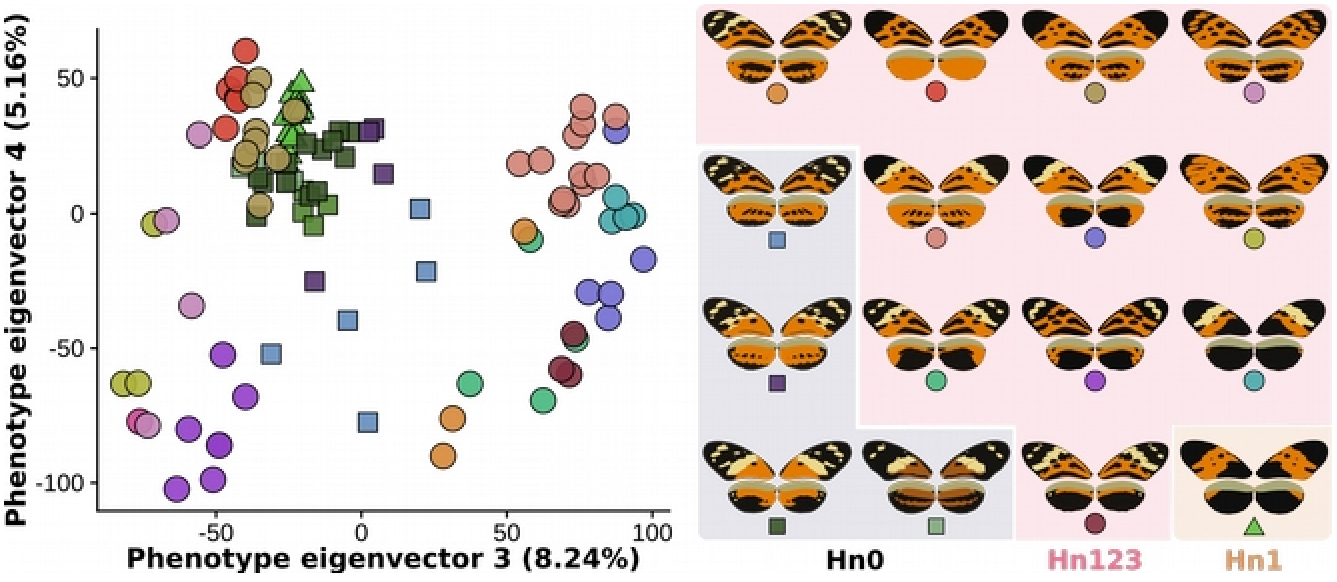


**Fig. S2 | Diversity of wing pattern in *H. numata*: components 3 and 4**

Principal component analysis of wing pattern variation observed across the *H. numata r*ange. Different morphs are depicted by different colors and different supergene genotype by different point shape (square : Hn0, circle : Hn123, triangle : Hn0).

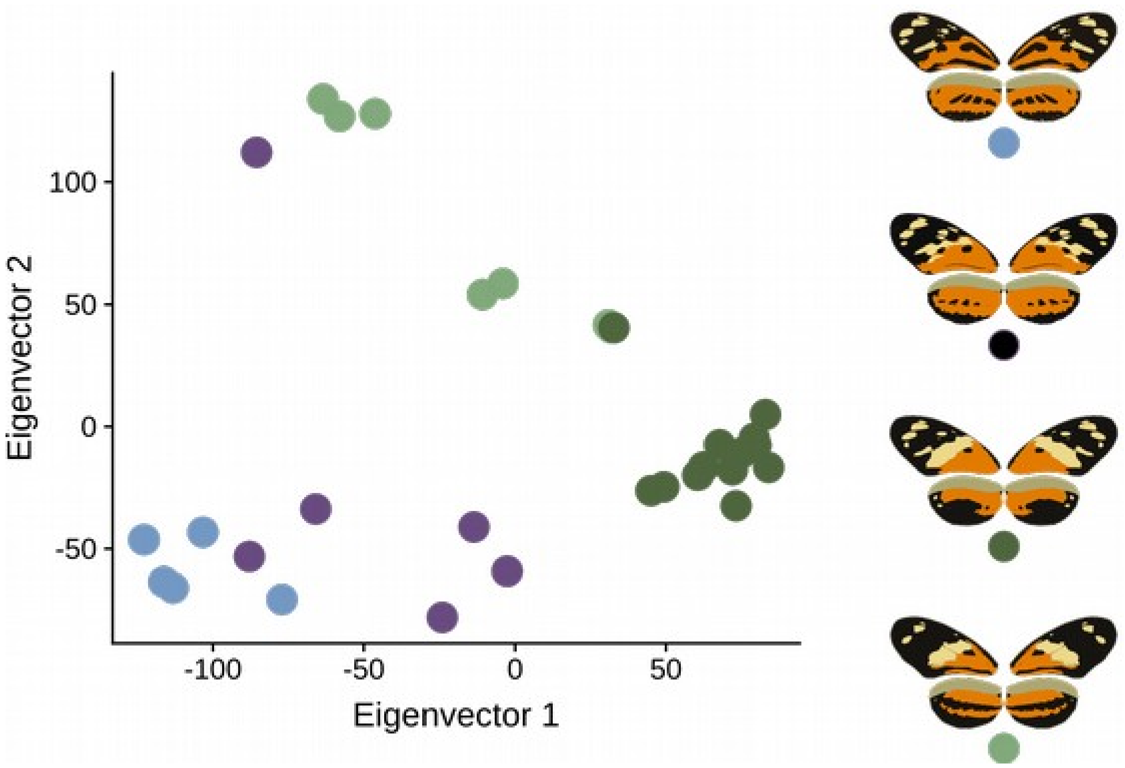


**Fig. S3| Diversity of wing pattern in Hn0 samples.**

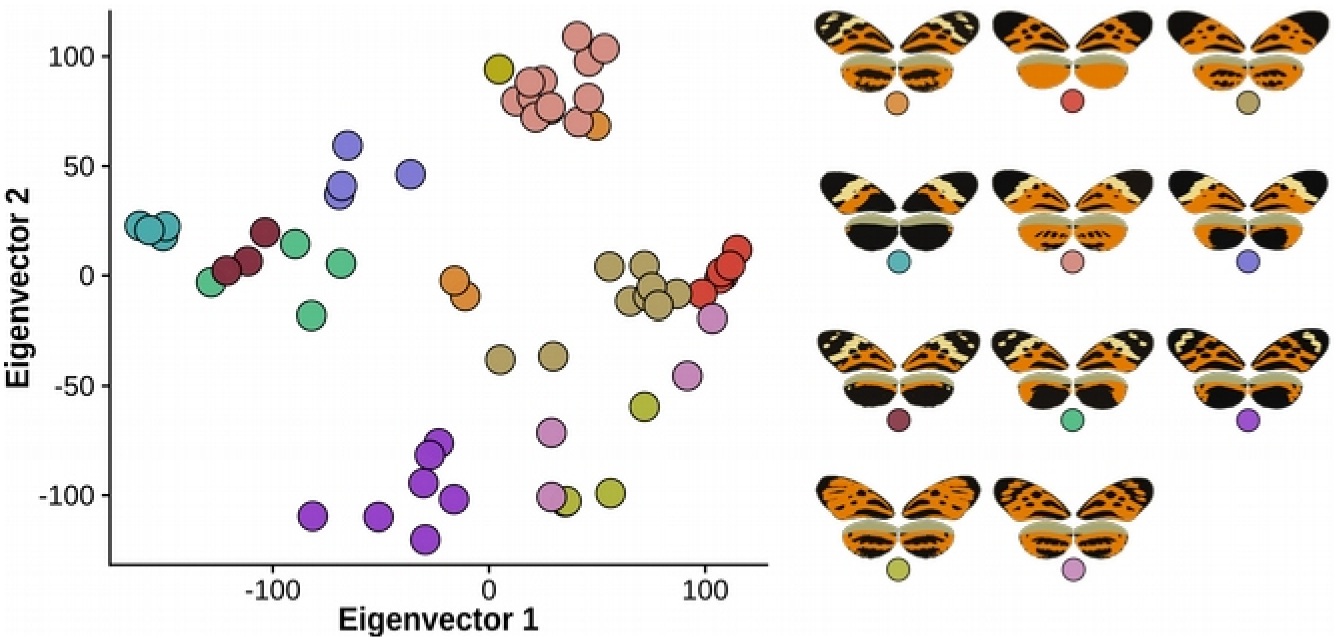


**Fig. S4 | Diversity of wing pattern in Hn123 samples.**

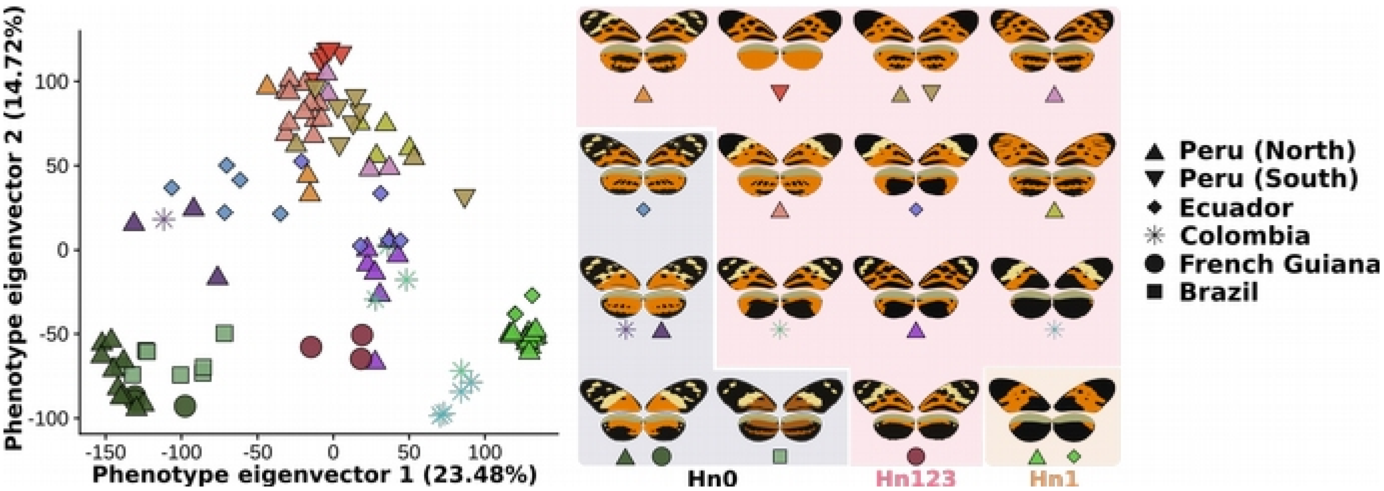


**Fig. S5 | The origin of sample is a poor descriptor of samples phenotype.**

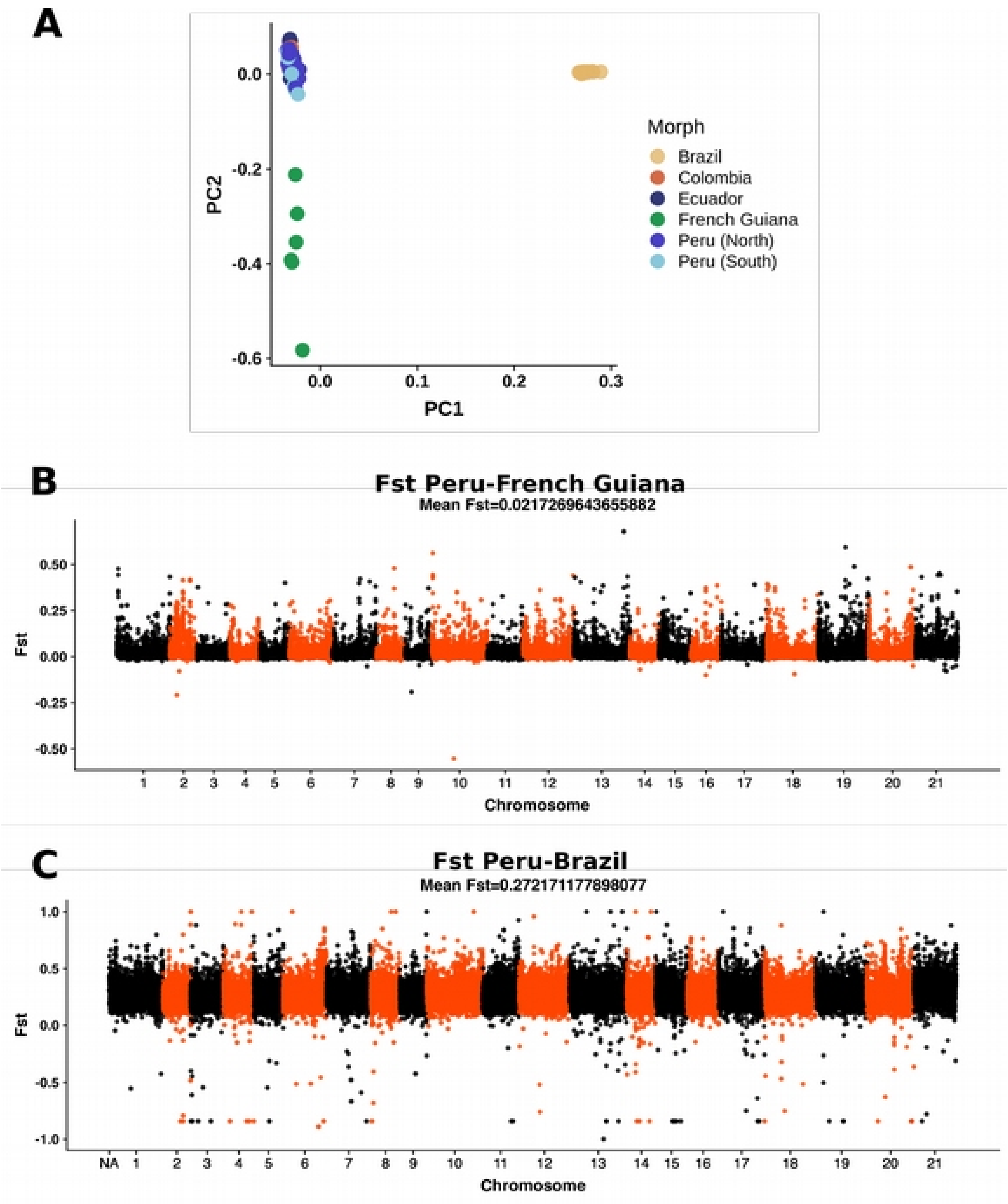


**Fig. S6 | Genetic structure in *H. numata*.**

**A** Principal component analysis displays three major clusters, formed by samples from Brazilian atlantic forest, by sample from French Guiana and by samples from Peru, Colombia, Ecuador and Colombia. **B** Fst analysis between Peruvian and Guianese samples. A low differentiation is observed between these two subsets of samples. **C** Fst analysis between Peruvian and Brazilian samples. A substantial differentiation is observed between these two subsets of samples.

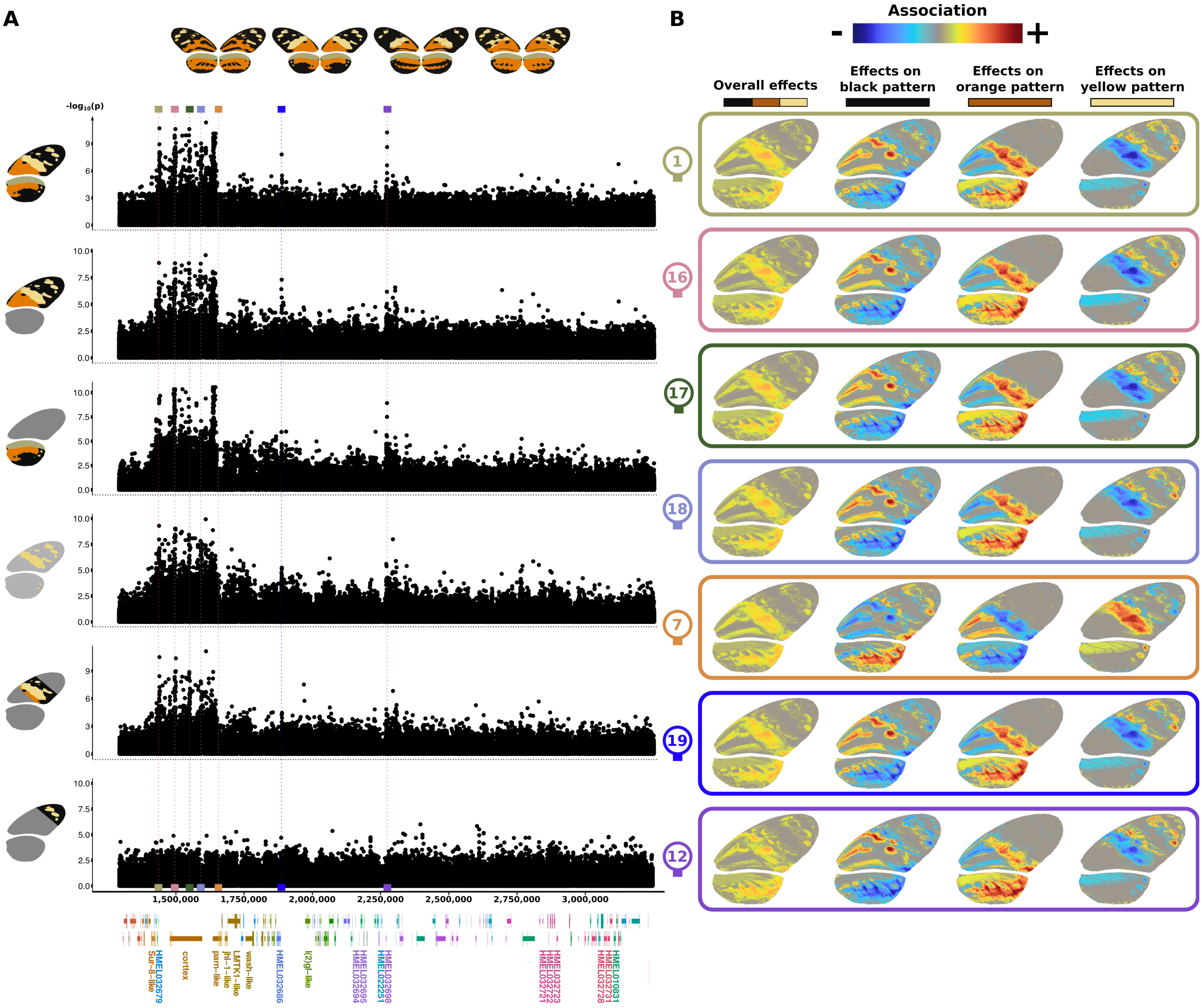


**Fig. S7 | Association study with Hn0 samples.**

**A** Multivariate association studies computed on Hn0 morphs and on different partitions of the wing: Hind and forewing together, forewing, hindwing, yellow pattern on both wings, middle part of forewing and tips of forewings. Coloured rectangles below plots indicate region displaying peak of association with one of the phenotype studied in Hn0. Data for Hn123 are presented in Fig. 3

**B** Phenotypic effect of each of the top SNPs from the 7 regions within the supergene with significant association with the wing pattern in Hn0 arrangement (colored rectangles). Overall effects are shown, as well as color-specific effects. Heatmaps from blue to red represent the strength of association of the derived allele for every pixel, with gray meaning no effect, blue negative association and red positive association. For Hn0 samples, all SNP have very similar phenotypic effect, since the reduced phenotypic diversity and sampling do not allow todecorrelate the distinct elements of the phenotype.

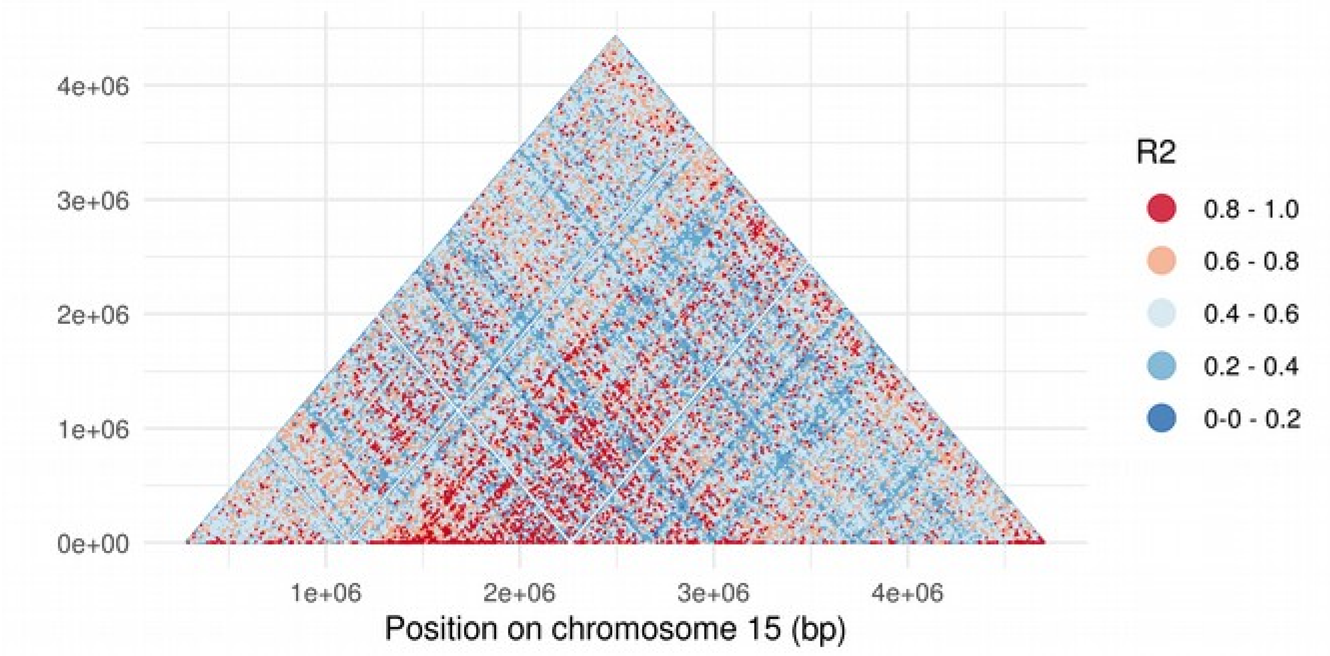


**Fig. S8 | Linkage disequilibrium heat map for Hn123 samples.**

Only samples homozygous for the three inversions are used in this analysis.

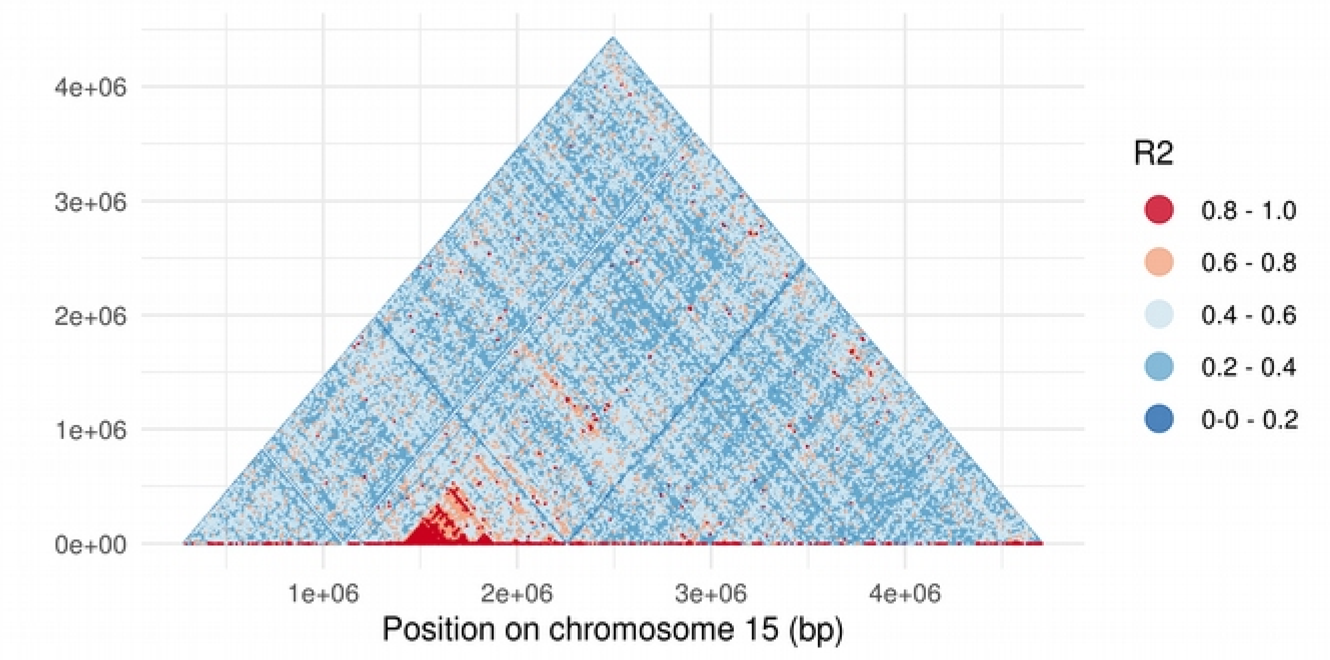


**Fig. S9 | Linkage disequilibrium heat map for Hn0 samples.**

Only samples homozygous for the ancestral gene order (Hn0) are used in this analysis.

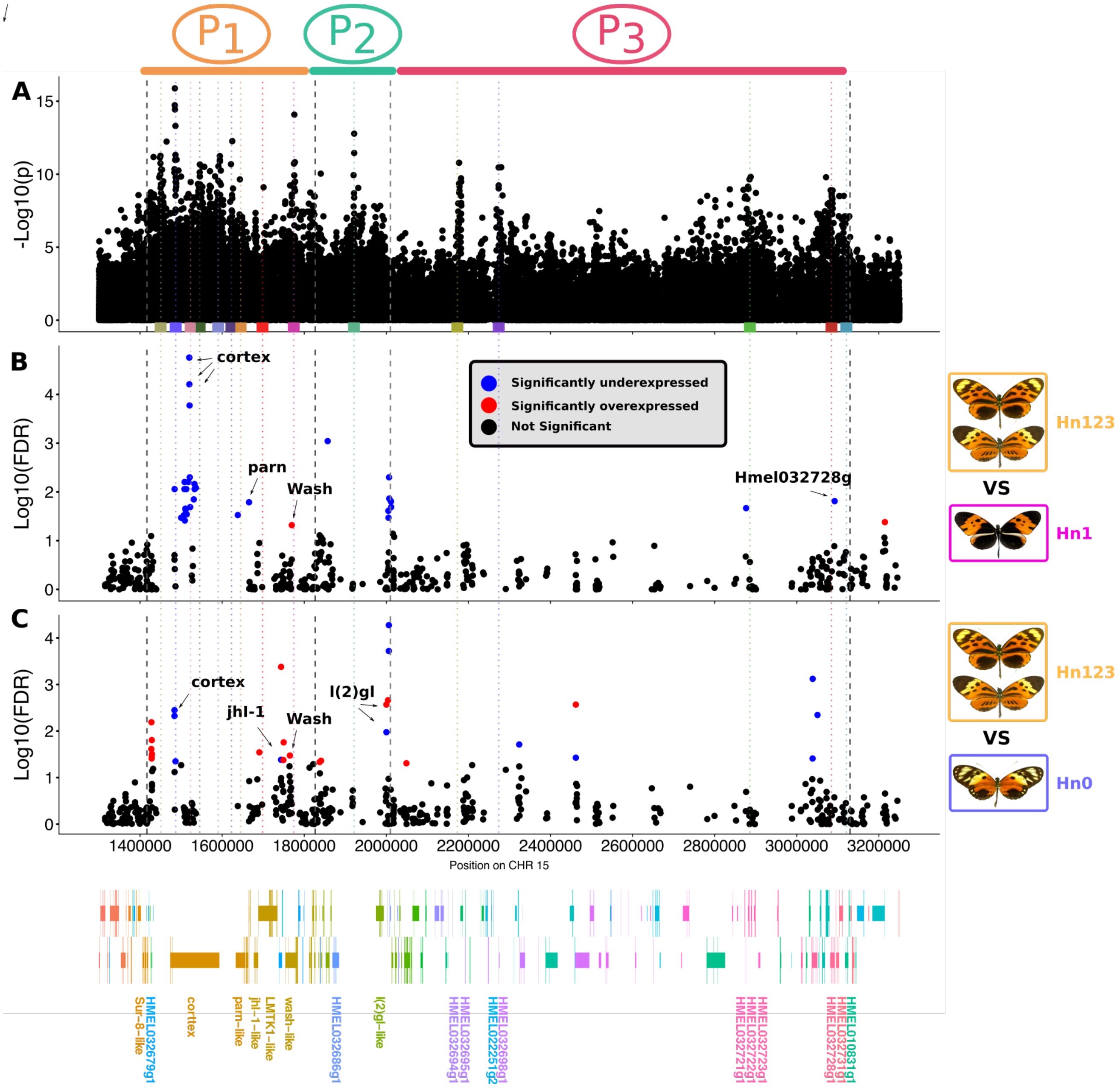


**Fig. S10 | Differential expression analysis at P supergene.**

**A,** Association computed on Hn123 samples. Similar to Fig. 3. The position of the peak of association found in either Hn123 are indicated by colored rectangles below the plot. **B,** Differential expression analysis comparing the RNA expression in early pupal (24h) wings discs between Hn123 and Hn1 supergene allelic classes. **C,** Differential expression analysis comparing the RNA expression in early pupal (24h) wings discs between Hn123 and Hn0 supergene allelic classes. For Hn123, two morphs were merged to benefit from increased statistical power. The -log10 of the false discovery rate is plotted along the chromosome 15, with each dot representing a different transcript. The *H. melpomene* gene position is represented bellow the plot.

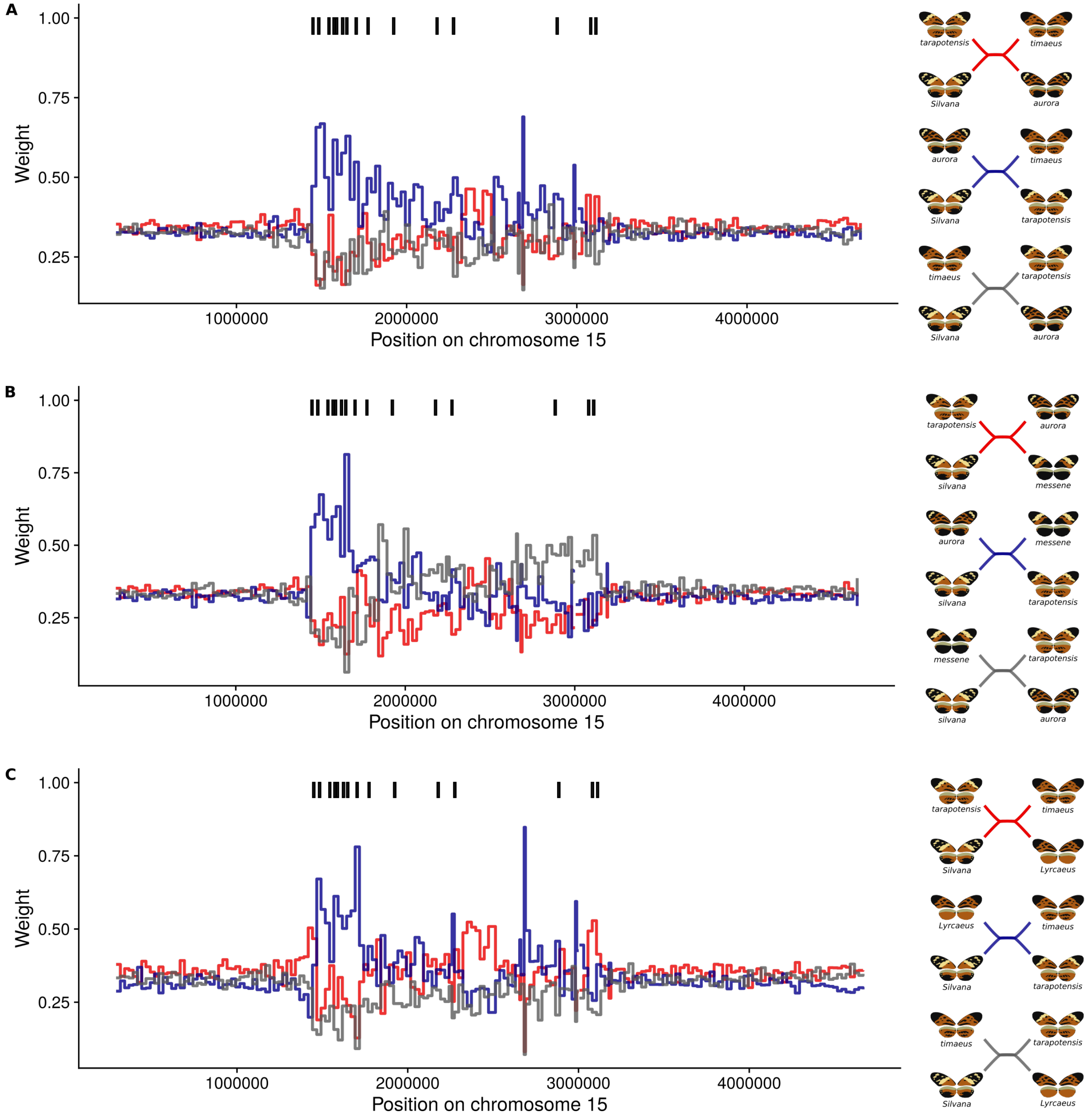


**Fig. S11 | Phylogenetic relationship of H. numata morphes at the supergene.**

The weight attributed to each topology was calculated using twisst (see methods). Squares above the plot indicates regions associated with wing pattern variation in Hn123 (coloured in other figures; Table S2) **A.** Relationship between *silvana* (outgroup), *aurora*, *tarapotensis* and *timaeus*. **B.** Relationship between *silvana* (outgroup), *messene*, *tarapotensis* and *aurora*. **C.** Relationship between *silvana* (outgroup), *lyrcaeus*, *tarapotensis* and *timaeus*.

**Table S1 | List of samples and sequencing statistics (provided separately)**

**Table S2 | List of most association regions and variants (provided separately).**

We defined here a region of association as a 10 kb region displaying a clear peak of association (several SNP with $p < 10^{-7}$). The choice of region size is arbitrary and a finer or larger region size can alter the results. Regions associated with wing-pattern variation in both Hn123 and Hn0 allelic classes are highlighted in yellow.
